# AI-based separation of malignant cell- and microenvironment-specific gene expression from bulk RNA sequencing enhances biomarker interpretation

**DOI:** 10.64898/2026.04.28.721082

**Authors:** Anastasia Zolotar, Daniil Wiebe, Arman Petrosyants, Boris Shpak, Natalia Khotkina, Valentina Beliaeva, Ekaterina Ivleva, Linda Balabanian, Maksim Chelushkin, Daniiar Dyikanov, Maria Savchenko, Sheila T. Yong, Danil Litvinov, Anastasia Zotova, Alexander Kuznetsov, Aleksandr Zaitsev, Anna Sharun, Artem Kosmin, Krystle Nomie, Mary Abdou, Aleksandr Sarachakov, Alexander Bagaev

**Affiliations:** BostonGene Corporation, Waltham, MA, USA; Applied AI Institute, Moscow, Russian Federation

## Abstract

Bulk RNA sequencing (RNA-seq)-based gene expression analysis is a promising tool for personalized cancer diagnostics, disease monitoring, and treatment decision-making. However, its clinical utility is limited by interference from non-malignant tumor microenvironment cells, which can dominate transcript data in low-purity tumors. While cell deconvolution methods like Kassandra can predict digital cell percentages from bulk RNA-seq, approaches for delineating the gene expression contribution of tumor compartments remain limited. To overcome this limitation, we developed Helenus, a machine-learning-based tool that separates gene expression between malignant and non-malignant cells. Trained on over 200 million synthetic RNA profiles representing diverse tumor types and purities, Helenus demonstrated high accuracy in separating gene expression origin. Helenus also uncovered true genomic-RNA correlations such as copy number alterations and the expression of therapeutic antibody-drug conjugate targets specifically on tumor cells. Helenus provides critical insights into tumor biology and immunotherapy response by precisely identifying biomarker expressions, paving the way for more effective personalized cancer care.

**Significance:** Helenus extracts gene expression profiles of cancerous and non-cancerous compartments of tumor biopsies from bulk RNA-seq data, enabling the determination of how the expression of specific genes affects malignancy and tumor immunity.

## INTRODUCTION

In clinical diagnostics, biomarker expression is often evaluated using immunohistochemistry (IHC), a method that analyzes only a small number of genes per test, restricting the scope of biomarker discovery (1). A powerful tool for whole-transcriptome analysis of tumors, bulk RNA sequencing (RNA-seq) can overcome this limitation because of its high-throughput nature that enables the discovery of a wide range of clinically relevant biomarkers.

However, a key challenge in adopting bulk RNA-seq for clinical practice is the heterogeneous composition of tumor biopsies, which include both malignant cells (MCs) and tumor microenvironment (TME) cells. In many cases, TME cells dominate the transcriptomic output, masking MC-specific gene expression and complicating biomarker interpretation. This can lead to confounding results, as TME-derived transcripts may overshadow the expression of critical biomarkers in MCs.

Overcoming this challenge requires approaches that can extract TME- and MC-specific gene expression profiles from bulk RNA-seq data so that the contribution of specific genes to the malignancy can be properly ascertained. Accurately identifying and quantifying biomarkers in specific tumor compartments is essential for predicting tumor progression, assessing treatment efficacy, and advancing drug development (2,3). Therapeutic targets expressed on tumor cells such as TROP2, FOLR1 (4), and HER2 play a critical role in the efficacy of monoclonal antibody therapies for ovarian, urothelial, and breast cancers (5). Similarly, prognostic markers like BAP1, which is often lost in malignant tissues and associated with poor outcomes, pose challenges due to their high expression in normal cells, making tumor purity a confounding factor (6). Moreover, benign stromal and immune cells in the TME can influence tumorigenesis through bidirectional signaling with MCs, as seen with biomarkers like LAG3, TIGIT, and PD-L1 (7,8).

The heterogeneous cellular composition of tumors highlights the need for robust methods that unmask the gene expression profiles of select tumor components. Current methods such as quantitative reverse transcription PCR (qRT-PCR) are highly sensitive but limited to analyzing specific biomarkers, while IHC provides spatial resolution but lacks scalability. RNA-seq offers the advantage of simultaneously analyzing a large number of genes (1,9). However, existing RNA-seq-based methods, such as purity correction (10) and linear subtraction, oversimplify tumor heterogeneity, resulting in imprecise analyses. Machine learning (ML)-based deconvolution tools, such as BayesPrism (11) and CIBERSORTx (12), rely on single-cell RNA-seq (scRNA-seq) reference data or predefined signature matrices. These methods are susceptible to batch effects, tissue quality limitations, and biases introduced during cell isolation (11,13).

We previously developed Kassandra, an ML-based cellular deconvolution algorithm trained on artificial transcriptomes that can accurately identify up to 51 cell populations in bulk RNA-seq data from tumors and blood (14). Having accomplished the critical task of separating cell populations in tumors and blood, we now introduce Helenus, a complementary ML algorithm that separates MC and TME gene expression profiles solely from bulk RNA-seq data.

Helenus has demonstrated exceptional accuracy in reconstructing the expression of 1,078 clinically relevant genes in MCs and the TME, including those encoding oncogenes, oncosuppressors, and immune response genes, as well as multiple gene signatures. Using cell population predictions from Kassandra, Helenus subtracts the weighted average expression of the TME and adjusts for tumor purity to provide robust analyses of gene expression within tumor samples. Through computational and experimental validation, Helenus has shown high performance in separating MC and TME gene expression profiles compared to existing approaches, which may translate to enhanced clinical interpretability of biomarker expressions from bulk RNA-seq. Helenus uncovered TME- and MC-specific expression of clinically relevant therapeutic antibody-drug conjugate (ADC) targets, showing strong concordance with IHC, and demonstrated enhanced associations of biomarkers with prognosis and therapy responses.

## RESULTS

### ML-driven separation of malignant and TME gene expression using simulated transcriptomes

Computational framework Helenus was designed to accurately differentiate gene expression profiles of MCs and TME cells using bulk RNA-seq data (**Fig. 1A**). ML tools like Helenus require extensive training datasets to capture the broad variability in tumor cellular phenotypes. However, existing tumor gene expression data are limited and fragmented across open-source databases. To address this, we simulated tumor transcriptomes by computationally mixing RNA-seq data from purified MCs and various TME cell types, leveraging a previously established approach for reconstructing tumor heterogeneity (14).

**Figure 1.**
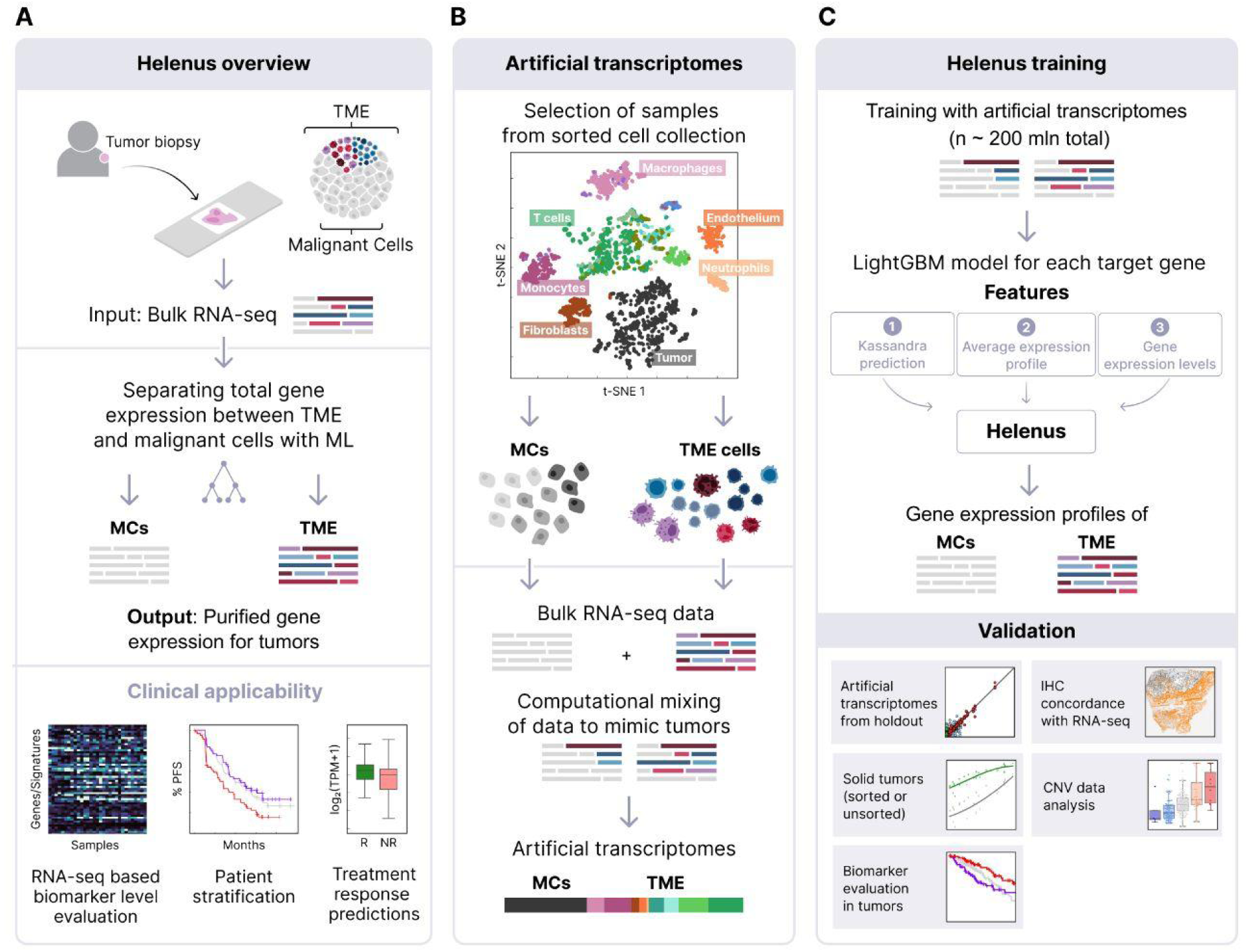
The construction of Helenus and its applicability. **(A)** Helenus reconstructs MC and TME expression profiles of tumor samples. The clinical utility of Helenus includes biomarker evaluation and prediction of prognosis and treatment response. **(B)** The training dataset consisted of ∼200 million artificial transcriptomes generated by computationally mixing RNA-seq data from sorted solid tumor and TME cells, retrieved from Kassandra’s collection. **(C)** (Top panel) Each of the 1,078 models was trained on RNA-seq data from the artificial transcriptomes (top) with three sets of features to generate the expression profiles of TME cells as the output. Consequently, the whole expression profile consisted of 1,078 target genes, each characterized by an individual model. (Bottom panel) Expression analysis by Helenus was tested with computational and experimental data in multiple validation approaches to reconstruct the MC and TME expression profiles.

The training dataset included 1,496 samples from 177 distinct solid tumor cell lines representing 19 solid tumor types and 4,575 TME samples comprising 137 unique TME cell types from the Kassandra database (**Fig. 1B**). To generate a comprehensive dataset, we created 200,000 artificial transcriptomes for each gene to mimic diverse tumor types and purities, culminating in ∼200 million simulated transcriptomes (**Fig. S1A**).

The Helenus models were trained to predict the gene expression contributions of TME cells within a tumor sample, as the inherently unstable gene expression patterns in MCs—driven by mutations, chromosomal instability, epigenetic alterations, and dysfunction in regulatory and transcriptional processes—render them unsuitable prediction targets. As such, pure TME gene expression profiles served as reference targets. Initially, 1,885 gradient-boosting LightGBM models (15) were trained and tested, one for each target gene. Among these, 1,078 gene models passed our stringent selection criteria (**see Methods, Fig. S1B**) and were used to reconstruct the complete TME expression profile predicted by Helenus. The prediction of TME gene expression was informed based on three key features:

1. The proportion of TME cell types in the sample, as determined by Kassandra deconvolution (14).
2. The weighted average expression of the target gene in TME populations (transcripts per million [TPM]), derived from 4,575 TME samples in the Kassandra database.
3. Expression levels (TPM) of a curated set of 600–603 genes specific to TME and MC populations, selected from an initial pool of over 20,000 genes (**Fig. 1, S1C**).

Helenus then estimated MC gene expression profiles by subtracting TME contributions from the uncorrected total tumor expression levels. These profiles were further adjusted for tumor purity (**see Methods**).

We validated Helenus through multiple approaches and assessed its utility in tumor biopsy analyses to demonstrate its potential for biomarker-driven patient stratification and improved prediction of clinical outcomes (**Fig. 1C**).

### Helenus accurately reconstructed tumor and microenvironment gene expression profiles in artificial tumor transcriptomes

To validate Helenus, we used a holdout test dataset of bulk RNA-seq samples representing TME cell populations (100 stromal samples and 209 peripheral blood mononuclear cell (PBMC) samples) and 50 solid cancer cell line samples. These samples spanned five cancer types—lung adenocarcinoma (LUAD), prostate cancer (PRAD), pancreatic ductal adenocarcinoma (PAAD), cutaneous melanoma (SKCM), and breast invasive carcinoma (BRCA)—with tumor purity levels ranging from 0% to 100%, based on MC RNA fractions. Although predictions for MC gene expressions by Helenus were closer to the reference MC profile than bulk RNA-seq for samples with any purity, a limit of detection (LOD) of >20% MC RNA was set for subsequent analyses, as lower tumor purity levels are below clinically acceptable thresholds (**Fig. S2A**).

The reconstructed gene expression profiles of TME and MCs were compared against reference profiles derived from pure TME RNA and MC RNA populations, as well as the artificial transcriptomes before Helenus reconstruction (**Fig. 2A, top**). Helenus successfully reconstructed TME and MC profiles that closely matched their respective references, unlike the initial artificial transcriptomes that showed marked discrepancies. TME reference profiles incorporated mixtures of PBMCs and stromal cells (e.g., fibroblasts, endothelial cells) in varying proportions, as determined by Kassandra deconvolution (**Fig. 2A, bottom**). Despite these complex mixtures, Helenus accurately reconstructed gene expression profiles of stromal and immune cells within the TME.

**Figure 2.**
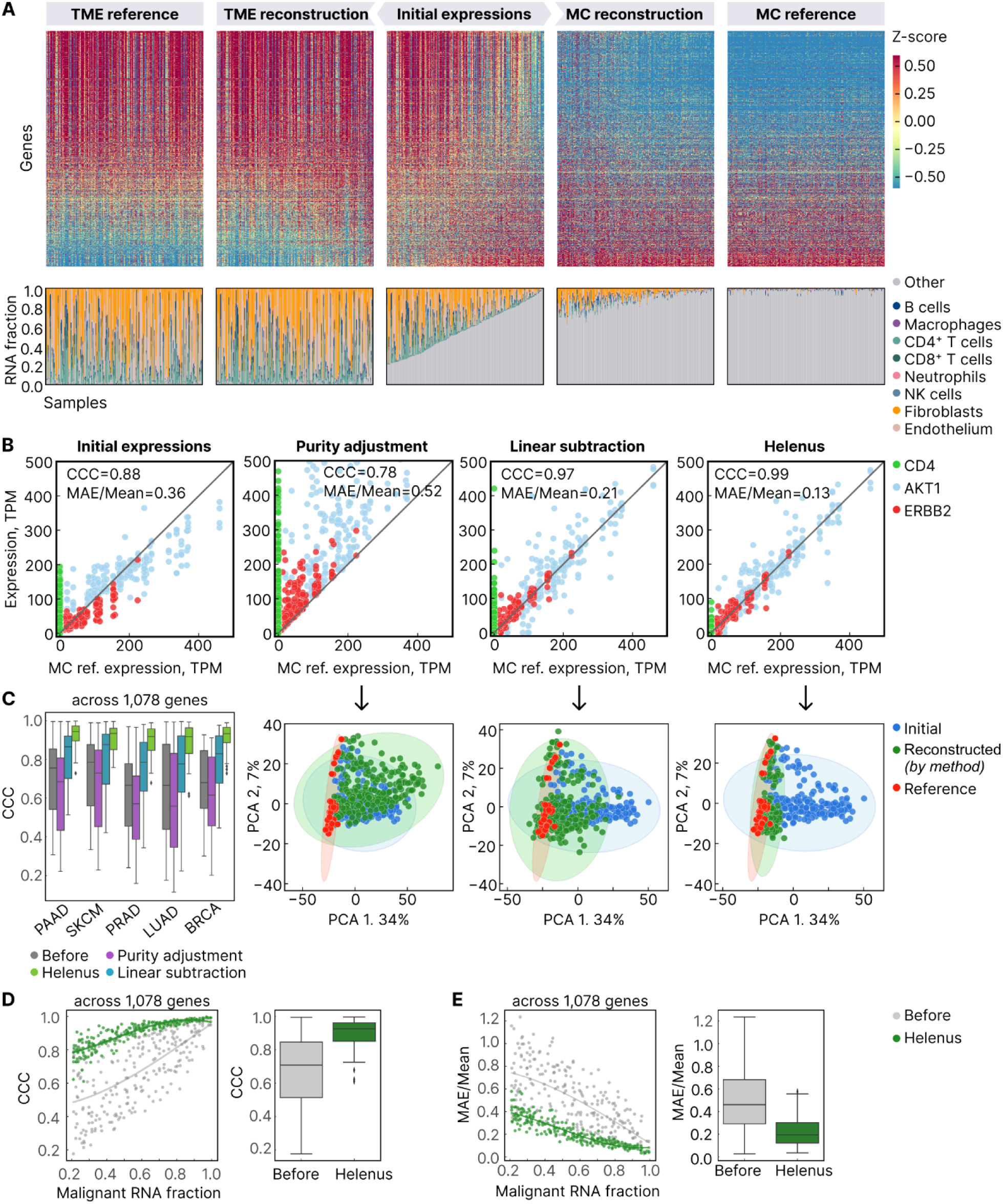
Gene expression origin separation in the artificial transcriptomes from holdout samples using Helenus. **(A)** Heatmap of TME and MC expression profiles reconstructed by Helenus as compared with their respective references and initial expression profiles (before Helenus). Datasets for the five cancer diagnoses analyzed were merged together and transformed via standardization scaling. The genes (rows) follow an ascending order based on the difference between mean expression level in TME and MC references. The barplots show the Kassandra-predicted RNA fractions in the TME (both immune and stromal cells). The ‘Other’ grouping is expected to predominantly include MCs. **(B)** Comparison of the expression profiles of three selected genes before and after the specified reconstruction across all five diagnoses. Each dot represents a sample. **(C)** Pairwise CCC of all 1,078 genes comparing the quality of expression level prediction in the MC fraction by the three methods for five cancer types (left). The principal component analysis (PCA) projections for the expressions of 1,078 genes visually demonstrate their distances before (blue) and after each reconstruction method (green), compared to reference expressions in MCs (red). The percentage (%) of the variance explained per PCA component is shown: initial expressions, the most diverse, were input first; then, the MC reference expressions which differed the most from initial expressions were added; finally, reconstructed expressions were projected to compare their closeness to initial and reference points. PCA projections for each diagnosis are shown in **Fig. S2C**. **(D)** CCC and **(E)** MAE/Mean pre- and post-application of Helenus for samples with varying MC purities (MC RNA fraction ranging from 0.2 to 1.0) across all five diagnoses. Each dot represents the average metric value for all 1,078 genes for a given sample (n=246 artificial transcriptomes). Box plots in C, D, and E show median and the 25th and 75th percentiles. Whiskers indicate the farthest datapoint within 1.5 * IQR (interquartile range).

We then compared Helenus’s performance against the performance of the following alternatives:

1. **Purity adjustment alone**: This method corrects gene expression levels by dividing the total expression by the estimated MC RNA fraction (10).
2. **Linear decomposition:** These methods consider the total tumor expression as a linear combination of MC- and TME-based gene expression. In the simplest form, they subtract a weighted average TME-based gene expression from bulk gene expression to reconstruct MC-specific profiles (CIBERSORTx) (12).

To illustrate the accuracy of these methods, we evaluated them across five cancer types using three clinically relevant biomarker genes with distinct expression patterns (as defined by their expression in sorted cells and cancer cell lines used for training): *CD4* (TME-specific, e.g., in CD4+ T cells), *ERBB2* (MC-specific), and *AKT1* (expressed in both MCs and TME cells). We used Mean Absolute Error/Mean (MAE/Mean) to normalize the absolute errors, thus enabling direct comparisons across genes with different expression magnitudes. In addition, we computed the concordance correlation coefficient (CCC) to measure how closely each reconstructed MC gene expression profile matched the corresponding reference MC profile (**see Methods**). Linear subtraction outperformed purity adjustment alone, reducing MAE/Mean from 0.36–0.52 to 0.21 and increasing CCC from 0.78–0.88 to 0.97 (**Fig. 2B**).

Unlike these approaches, Helenus predicts gene expression contributions using trained models and applies purity adjustment to renormalize expression levels (**see Methods**). Importantly, Helenus surpassed both methods, achieving a reduction in MAE/Mean to 0.13 and increasing CCC to 0.99 for the three biomarker genes (**Fig. 2B**). Helenus consistently demonstrated lower MAE/Mean and higher CCC across all 1,078 target genes regardless of gene expression levels (low, medium, or high; **Fig. 2C, Fig. S2B**) and outperformed the linear reconstruction methods (purity adjustment and linear subtraction). Principal component analysis (PCA) revealed that while linear subtraction reduced the distance between reconstructed and reference MC profiles, Helenus’s predictions were the most accurate, closely aligning with the reference profiles (**Fig. 2C, Fig. S2C**). In contrast, purity adjustment increased the divergence from reference profiles, highlighting its limitations. Moreover, Helenus accurately reconstructed MC gene expression profiles across a wide range of tumor purities in the artificial transcriptomes (**Fig. 2D–E, Fig. S2A**). Application of Helenus improved the median CCC from 0.71 to 0.93, while MAE/Mean decreased from 0.46 to 0.19 (**Fig. 2D–E**). These results demonstrate Helenus’s superior performance in reconstructing MC gene expression profiles compared to existing methods.

### Helenus separated expression of *TROP2* and *FOLR1* in malignant and TME cells and exhibited strong concordance with IHC

Having established Helenus’s performance in separating TME and MC expression profiles, we evaluated whether the predicted gene expression levels are comparable to biomarker quantification obtained via immunohistochemistry (IHC), the current standard in clinical practice, for two ADC therapeutic targets: TROP2 (*TACSTD2*) and folate receptor alpha (*FOLR1*) (**Fig. 3A**).

**Figure 3:**
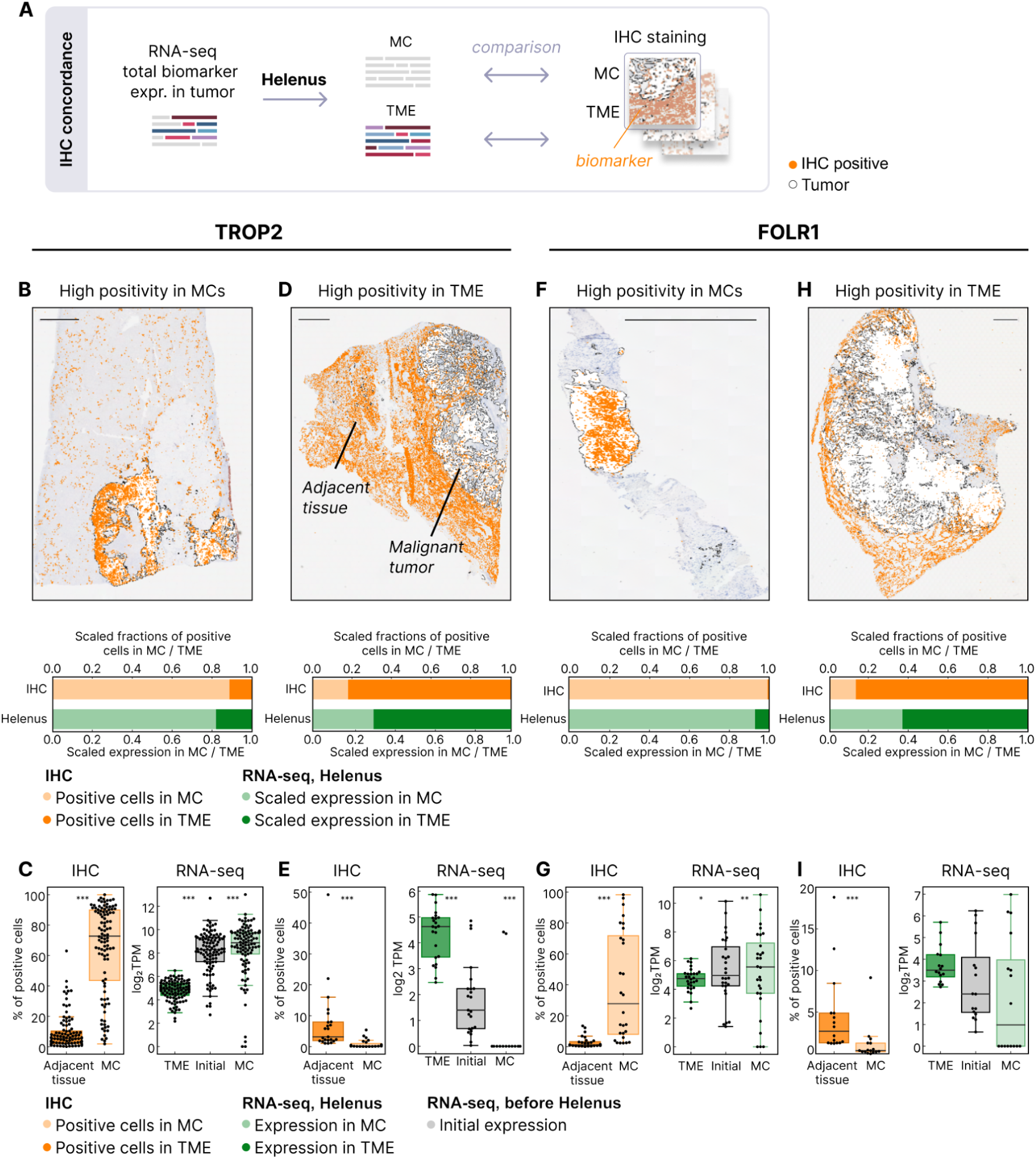
Concordance between Helenus-predicted RNA expression levels and IHC scores for TROP2 and FOLR1 in tumors. Samples were categorized into two groups based on the predominant location of positive staining, either within the TME or in MCs. Concordance between Helenus reconstructions and IHC staining was assessed. **(A)** Schematic depicting comparison of biomarker levels as assessed by IHC and Helenus-corrected RNA-seq. **(B)** Image of a representative sample with predominantly TROP2-stained MCs. **(C)** TROP2 expression based on IHC compared to RNA-seq before and after Helenus (n = 102). **(D)** Image of a representative sectioned biopsy sample stained for TROP2 with a majority of TROP2-positive cells distributed within the TME. The barplot indicates the abundance of TROP2-positive cells, and the ratio of *TROP2* expression levels in MCs relative to the TME. **(E)** Boxplots showing the percentages of positively stained cells for TROP2 among MCs and in the TME (left), and the initial *TROP2* expression (before Helenus) from RNA-seq compared to Helenus-reconstructed expression profiles of MCs and the TME (n = 23 samples). **(F)** A representative sample with MC-dominant distribution of FOLR1 protein. **(G)** FOLR1 expression based on IHC versus RNA-seq before and after Helenus (n = 28). **(H)** Slide and barplot for a representative sample exhibiting a TME-dominant FOLR1 staining. **(I)** Boxplots showing protein expression and Helenus-reconstructed RNA expressions in TME and MCs, as well as initial *FOLR1* expression before Helenus (n = 16) Boxplots indicate median and the 25th and 75th percentiles, with whiskers extending to the lowest and highest value within 1.5x of the interquartile range. Wilcoxon signed-rank test compared the difference between IHC scores of the TME and MCs (boxplots on left), and between initial RNA expressions and those of the TME or MCs (boxplots on right). Significant *P* values are indicated on the boxplots: *: *P*<0.05, **: *P*<0.01, ***: *P*<0.001. Scale bars in the top images represent 2 mm.

We categorized IHC-stained samples into two groups based on staining patterns. **Group 1** contained samples with a higher percentage of positively stained MCs compared to adjacent TME cells (examples in **Fig. 3B, F**), while **Group 2** contained samples with high IHC staining in adjacent TME cells but not in MCs (examples in **Fig. 3D, H**). Helenus’s reconstructed MC expression profiles revealed high *TROP2* RNA expression in MCs for Group 1, consistent with high TROP2 IHC positivity in MCs but not in adjacent tissues (**Fig. 3C**). Conversely, Helenus identified elevated *TROP2* expression in the TME for Group 2, where IHC showed low or absent TROP2 staining in MCs but high positivity in adjacent TME cells (**Fig. 3E**). Similarly, Helenus demonstrated strong concordance with IHC readouts for FOLR1 expression levels across both groups (**Fig. 3F–I**). Importantly, Helenus’s reconstructed expression profiles for *TROP2* and *FOLR1* were significantly different from unadjusted bulk RNA-seq levels but instead closely matched observed IHC scores. Benchmarking Helenus against IHC, these findings demonstrate Helenus’s capability as a complement to IHC, underscoring its precision in discerning biomarker expression and its cellular origins.

### Helenus refined interpretation of prognostic markers and therapy responses in tumors

Next, we benchmarked Helenus’s gene expression corrections against reference copy number variation (CNV) annotations in tumor specimens. Clinically actionable CNVs are typically associated with changes in gene expression levels (16). However, analysis of 2,204 samples with purity < 0.5 from TCGA revealed that uncorrected expression profiles did not correlate strongly with CNVs. Specifically, samples with complete gene loss (−2 CNV) often exhibited misleadingly high expression, and overall, gene expression levels showed weak correlation with CNVs (Spearman’s ρ=0.7, *P*=0.2; **Fig. 4A**).

**Figure 4:**
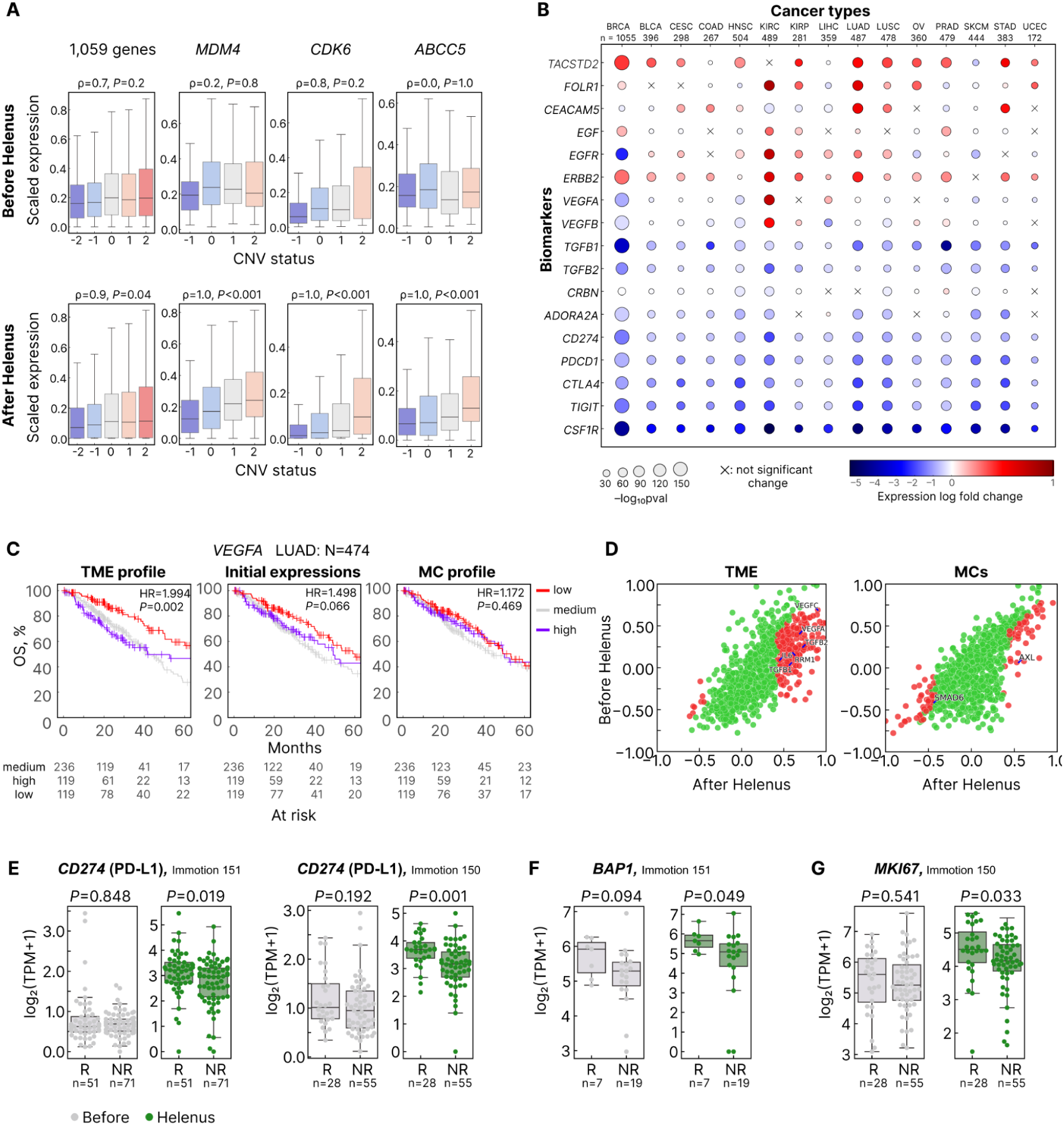
Corrected biomarker expression by Helenus and their relationship with patient survival and therapy response. **(A)** Boxplots showing scaled expressions in several diagnoses from TCGA cohorts, pre- and post-Helenus, for samples with different CNVs. Expressions were max scaled for each gene. Boxplots represent the median and 25th and 75th percentiles, with whiskers from the lowest and highest value within the 1.5x interquartile range. Gene copy number was determined from the CNA annotation of samples, considering ploidy. **(B)** Expression log fold change in TCGA cohorts post-Helenus. Log fold change > 0 refers to increased expression in Helenus-reconstructed biomarker expression in MCs compared to initial expression, while < 0 denotes decreased expression. Non-significant changes are marked with ‘X’ (Wilcoxon signed-rank test, Bonferroni correction). **(C)** Kaplan-Meier survival analysis of TCGA-LUAD based on *VEGFA* expression in the TME and MCs post-Helenus compared to initial uncorrected levels (logrank test). Patients were stratified by *VEGFA* expression levels: low (lower 25% quartile or 0q<x<0.25q), medium (0.25q<x<0.75q), and high expression groups (upper 25% quartile or 0.75q<x<1q). Hazard ratios (HRs) indicate the survival differences between high and low expression groups. **(D)** Cox proportional hazards regression coefficients for evaluating the association between survival of TCGA-LUAD patients and the expressions of 894 genes (circles). Coefficient of 1 or −1 indicates that survival depends on gene expression. *P* values in red represent values that showed statistical significance or were significant and decreased post-Helenus. *P* values in green represent values that remained statistically insignificant or increased post-Helenus (regardless of statistical significance). **(E, F, G)** Expressions of *CD274* and *MKI67* in the TME, and *BAP1* in MCs were compared between responder (R) and non-responder (NR) patients treated with combination atezolizumab/bevacizumab, before and after Helenus (Mann-Whitney U rank test). Primary site samples from IMmotion151 were selected with PD-L1 IHC scores < 1%, omitting liver metastases, to compose a more uniform dataset, while no sample exclusion was applied to the IMmotion150 dataset. The datasets were plotted separately to show reproducibility of analysis of *CD274* expression by Helenus. Individual sample data shown by circles; outliers by rhombuses.

In contrast, Helenus-corrected expression profiles demonstrated a strong and consistent dependence of gene expression on copy number, with negligible expression observed in samples with complete gene loss and a monotonic increase in expression corresponding to copy number amplification (ρ=0.9, *P*=0.04; **Fig. 4A**). For example, Helenus revealed positive correlations for *MDM4*, *CDK6*, and *ABCC5* (ρ*=*1*, P*<0.001), compared to the variable correlations observed in uncorrected profiles (ρ=0.2, 0.8, and 0, respectively; **Fig. 4A**). These findings validate the accuracy of Helenus’s expression purification in a real cohort of 2,204 patients.

Using bulk RNA-seq data from TCGA, we examined how Helenus-corrected expression levels altered the interpretation of therapeutic target expression. For markers typically overexpressed in MCs (e.g., *TACSTD2*/TROP2, *FOLR1*, *CEACAM5*, *EGF*, *EGFR*, and *ERBB2*/HER2), Helenus revealed higher expression in MCs compared to initial bulk RNA-seq levels. Similarly, Helenus highlighted higher expression levels of TME-specific markers (e.g., *ADORA2A*, *CD274*/PD-L1, *PDCD1*/PD-1, *CTLA4*, and *CSF1R*), consistent with their expected enrichment in the TME (**Fig. 4B**). These findings demonstrate that Helenus can accurately reconstruct expression profiles in concordance with known biology, improving the resolution of biomarker analysis.

By correcting expression values, Helenus can enhance the clinical interpretation of prognostic markers that may otherwise be confounded in bulk RNA-seq data. For instance, in TCGA-LUAD cohort, survival analysis revealed significant differences in overall survival (OS) for patients with high versus low expression of *VEGFA*, *VEGFC*, *TGFB1*, *TGFB2*, *IL6*, and *RRM1*, specifically in the TME (**Fig. 4C-D, Fig. S3**). Before correction by Helenus, these markers showed weak or non-significant associations with survival. Correction with Helenus revealed better OS for patients with low *VEGFA*/*VEGFC* expression in the TME compared to those with high expression (hazard ratios [HR]: 2.0 and 2.4, respectively; **Fig. 4C, Fig. S3**). These results align with the established pro-angiogenic role of VEGF secreted by tumor-infiltrating myeloid cells in the TME, exacerbating tumor progression (17–19). Similarly, Helenus-corrected MC profiles unveiled stronger correlations between MC-specific markers and survival outcomes. For instance, higher *AXL* expression correlated with worse OS (*P*=0.004; **Fig. 4D, Fig. S3**).

Helenus also improved the interpretation of therapy response biomarkers. Using clear cell renal cell carcinoma (ccRCC) samples from the IMmotion150 (EGAS00001002928) and 151 cohorts (EGAS00001004353) (20,21), we analyzed expression levels of *CD274*/PD-L1, *MKI67*, and *BAP1* (22–27), which were previously implicated in predicting response to atezolizumab/bevacizumab immunotherapy. Before correction by Helenus, expression levels of these markers did not differ significantly between responders and non-responders. After correction, however, higher *CD274* and *MKI67* expression in the TME and *BAP1* expression in MCs correlated strongly with therapy response (**Fig. 4E-G**).

## DISCUSSION

Transcriptome profiling can serve as a surrogate for multiple IHC stainings by measuring thousands of genes simultaneously, providing a comprehensive view of tumor biology in a single assay. While IHC and H&E staining remain the gold standards for determining biomarker positivity, they are limited in the number of markers they can reliably assess simultaneously. On the other hand, while bulk RNA-seq can address this limitation by profiling many genes in parallel, it only yields an average signal from a mixture of tumor cells and cells in the surrounding microenvironment. scRNA-seq offers cell-level resolution but is costlier and more complex. As such, these methods fall short in their utility in a clinical setting.

Methods that deconvolute bulk RNA-seq data into cell-type-specific signals can overcome this limitation and more closely approximate the level of detail provided by multiple IHC markers, making RNA-seq a viable, high-throughput alternative for clinical diagnostics and therapy selection (28). To address the shortcomings of RNA-seq and increase its clinical utility, we developed Helenus, a gene expression analysis tool that delineates the contribution of the TME from the total gene expression in a tumor and accurately reconstructs distinct gene expression profiles of MCs and the TME. The ML-based approach of Helenus accurately extracted gene expression profiles from bulk RNA-seq data for a panel of 1,078 genes, covering a large portion of clinically relevant biomarkers.

We showed signal unmixing to be correlated with spatial IHC resolution for markers TROP2 and FOLR1, demonstrating that Helenus predictions hold potential as complements for IHC in predicting spatial RNA expression in TME and MCs.

Our analysis revealed that bulk RNA-seq without correction often produced misleading correlations between gene expression and CNVs. For instance, uncorrected data showed weak or inconsistent associations, with high expression levels observed even in cases of complete gene loss. On the other hand, Helenus-corrected profiles aligned expression levels more closely with CNV data, producing robust and biologically accurate correlations. This capability of Helenus ensures that CNV-associated biomarkers are interpreted reliably in clinical and research settings.

While simple and straightforward, existing methods for analyzing gene expression profiles from tumor RNA-seq, such as purity adjustment and linear subtraction, often neglect the heterogeneity of samples, varying gene expressions, and dynamic regulatory patterns (29). The Helenus models are designed to account for these factors, leading to better performance. Unlike linear subtraction and purity correction, Helenus can elucidate complex and non-linear dependencies. To our knowledge, Helenus is the first algorithm that uses a gradient boosting framework (30) for tumor profile reconstruction.

Moreover, our previously developed approach for generating artificial transcriptomes (14) allowed us to create a large training dataset of ∼200 million mixtures, many more than what would be possible to collect from clinical patients. This is in concordance with recent advancements in AI where self-training models outperformed standard big data learning approaches (31). Unlike BayesPrism and CIBERSORTx, Helenus does not require pre-existing RNA-seq references of any kind, needing only bulk RNA-seq of samples to perform its function. This is an important advantage particularly in a clinical setting because regular users are unlikely to have access to such references, and obtaining them for simply analyzing a small number of patient samples would be a huge undertaking. Importantly, our strategy involving training gradient-boosted models on millions of artificial transcriptomes allowed Helenus to handle non-linearities and gene–gene interactions in a way that purely reference-based methods generally do not, enabling robust separation of malignant and TME-derived signals directly from bulk RNA-seq.

Helenus’s capability to analyze many genes at once yields a bird’s eye view on the gene expression landscape, facilitating analysis of multiple and complex biomarkers such as gene signatures. Precise measurements of biomarker expression are paramount to proper patient stratification to facilitate effective treatment. Here, we showed that Helenus-corrected profiles greatly improved the clinical utility of prognostic biomarkers, such as *VEGFA* and *VEGFC* in LUAD, as well as *CD274* (PD-L1), *MKI67*, and *BAP1* in ccRCC. These findings reinforce the utility of Helenus in translational medicine by more accurately identifying predictive biomarker expressions that guide personalized treatment decisions.

### Limitations

Helenus was developed primarily for solid, epithelial tumors. While normal epithelial cells may be present alongside MCs in some epithelial tumors and potentially confound gene expression deconvolution, we expect this effect to be minimal in well-dissected tumors and any biases in downstream analysis to be mitigated by comparison with normal tissues. Since its training data are largely composed of artificial transcriptomes modeling epithelial tumors, they do not represent malignancies dominated by non-epithelial cells (e.g., sarcoma) or purely hematologic cancers. Nonetheless, our validation showed that Helenus can be applied to some cancer types that are absent from the training dataset, like melanoma. Future development will focus on training Helenus with artificial transcriptomes recapitulating blood cancers, such as those of T cells for T-cell leukemia, B cells for follicular lymphoma, and myeloid cells for acute myeloid leukemia, to improve its performance on these cancer types.

A related constraint is potential performance degradation in tissues with unique cell compositions (e.g., liver, lymph nodes, or bone marrow) that were under-represented in the current training dataset. Addressing this will require expanded data sources, specialized training, and additional validation studies to capture the full diversity of these tissue environments.

Another limitation was the reduced performance observed in samples with tumor purity below 20%. While Helenus still improved gene expression detection compared to uncorrected bulk RNA-seq alone in these cases, future work will focus on enhancing model predictions for samples with low purity.

Lastly, Helenus presently reconstructs a selected panel of clinically relevant genes with gradient-boosting models, while the remaining genes were estimated via linear decomposition (the “Matrix M” method). A comprehensive expansion of gradient boosting-based reconstruction to all genes will demand larger open-source datasets, more refined artificial mixtures, and robust model optimization. These efforts, combined with a broader sampling of tumor and tissue types, will further expand the gene model repertoire and enhance Helenus’s clinical applicability.

## METHODS

### Sample collection for Helenus development

A total of 18,193 samples of sorted cells derived from tumor biopsies or blood and 715 cancer cell samples (both cell lines and sorted tumor cells) were collected from our database used previously for constructing the Kassandra cellular deconvolution tool (**Fig. 1**) (14). This database includes open-source RNA-seq datasets of sorted cells from ArrayExpress (32) and GEO (33). A total of 6,071 samples were selected for the construction of Helenus. The training dataset of artificial transcriptomes was obtained from 1,496 samples from 177 cancer lines across 19 solid tumor diagnoses and 4,575 TME samples. The TME samples consisted of 137 TME cell types from the Kassandra database: neutrophils, NK cells, macrophages, fibroblasts, endothelial cells, B cells, CD4^+^ T cells, CD8^+^ T cells, monocytes, and T helper cells, among others (14). All collected datasets included RNA-seq (read length >31 bp) without poly-A depletion and without the use of targeted panels. Several quality checks were performed, such as expression of cell-specific biomarkers (negative and positive biomarkers) and clusterization of similarly annotated samples. Samples with a total number of coding counts (of sequenced fragments) of less than 1 million were excluded.

### Generation of artificial transcriptomes for the training dataset

The collection of samples was used to create 200,000 artificial transcriptomes as training datasets for each model via computational mixing of RNA-seq data from the cancer cell samples and the TME samples. Samples from different cell types were randomly selected, averaged within the cell type, and summed into a final expression file in proportions resembling those in real tissues, as previously described (14). Since the number of available cell samples may vary significantly for different datasets and cell subpopulations, the number of samples were rebalanced per dataset and cell subtype to optimize the generation of artificial transcriptomes (see (14)). For selection of the MC proportion in each artificial transcriptome, fractions of tumor RNA were generated from a uniform distribution of 0 to 1. Consequently, the TME fraction was equal to *1 - tumor RNA fraction*. The RNA fractions of TME subpopulations followed a Dirichlet distribution. Further details have been published in Zaitsev et al. (2022) (14).

### Validation set of artificial transcriptomes from hold-out samples

Holdout samples for generating artificial transcriptomes for the first validation test comprised bulk RNA-seq data from TME cells sorted from both tissue (such as sorted stromal cells, e.g. fibroblasts, endothelial cells, n = 100) and blood (pure PBMCs, n = 209), as well as from cancer cell populations from five different cancer diagnoses (lung adenocarcinoma, prostate cancer, pancreatic ductal adenocarcinoma, cutaneous melanoma, and breast invasive carcinoma). Helenus was applied to reconstruct separate gene expression profiles for the MC and TME fractions for comparison with corresponding references. The references for reconstructed TME expression profiles were the profiles for pure sorted stromal cell and PBMC combinations. The pure MC references were profiles for the original cancer cell preparations. A limit of detection (LOD) threshold of 20% tumor purity (MC RNA fraction) was ascertained based on the performance of Helenus on analyzing artificial transcriptomes generated from holdout samples (**Fig S2A**).

### RNA-seq data processing

Bulk RNA-seq data were processed as previously described (14). Briefly:

1. Fastq files were processed using Kallisto (version 0.42.4 for Linux), followed by alignment to GENCODE transcriptome (version 23) and the GRCh38 human reference genome (excluding specific gene regions).
2. Single-end fastq files were processed with parameters -l 200 and -s 15.
3. Transcript-level expression was measured in TPM.
4. The same processing steps were applied to all cell-type datasets from GEO and ArrayExpress.
5. Quality control checks were performed using FastQC (v0.11.5 or later), FastQ Screen (v0.11.1 or later), and MultiQC (v1.4 or later).
6. Reference genomes for BWA aligner indices included genomes for multiple species and microbiome data from the NIH Human Microbiome Project, along with adapters and UniVec from NCBI.

### Gene expression data for Helenus

Helenus used TPM units to measure gene expression levels (34,35) and reduce batch effects between different datasets (14). Short RNA transcripts may strongly skew the detected expression distribution for a target gene (in TPM). As such, transcript filtering was applied to reduce variability in total expression. Non-coding RNAs as well as short transcripts of T-cell receptor (TCR)- and B-cell receptor (BCR)-coding genes, histone-coding genes, mitochondrial genes, and 48 additional transcripts were excluded from expression values, as explained in Zaitsev et al. (2022) (14).

### General structure of Helenus

The input for Helenus is bulk RNA-seq data of a sample for the expression of at least 18,792 genes. The gene expression counts of the sample are transformed to TPM and the summed expression of 18,792 genes is normalized to 1 million TPM. Firstly, the core Helenus prediction using a gradient boosting LightGBM model determines the TME contribution to the total expression profile of the tumor. Secondly, a separate LightGBM gradient boosting model predicts the RNA fraction of the TME based on the expression level of 1,305 genes used to train the tumor purity prediction model (**Fig. 1**). It is assumed that the gene expression profile of a sample is the sum of gene expression in MCs and TME.

Consequently, the fraction of malignant RNA is estimated as:

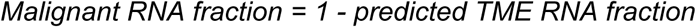

The core Helenus prediction and the purity prediction model are used together to estimate the TPM-renormalized, reconstructed gene expression profile of pure TME and MC compartments, respectively, according to the following formulas:

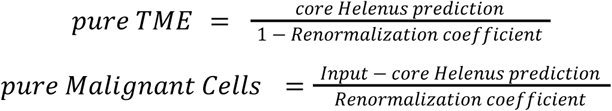

where the core Helenus prediction is gene expression in the TME. The *renormalization coefficient (x)* is derived from the purity prediction model as *1 - output of our purity prediction model = 1 - TME fraction*, or succinctly, *the MC RNA fraction*. TPM renormalization recovers the summed expression of all genes back to 1 million TPM after subtraction of the TME contribution. Overall, the gene expression in the sample is defined as follows:

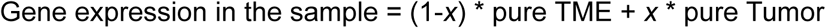

where *x* is the MC RNA fraction, ‘pure TME’ refers to gene expression in the TME only, and ‘pure Tumor’ refers to the expression in the tumor only, where summed expression of all genes equals 1 million TPM.

### Core Henenus prediction

The output of each LightGBM model for the core Helenus prediction is defined by *(1-x) * pure TME*, or weighted contribution of the TME component, where *x* corresponds to the malignant tumor RNA fraction. We indirectly estimated the MC expression profiles by predicting the expression profile of the TME, which is relatively stable compared to the tumor genome.

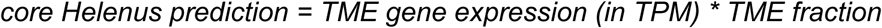

Gene expression in the TME is the result of a linear combination of individual cell expression profiles within that tissue (14):

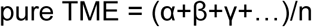

where [α, β, γ…] each represents the expression profile of respective cell types and n is the length of the vector.

Each core Helenus prediction model estimates the expression of one target gene independently from one another. Each gradient boosting model was trained with three sets of features to predict TME expression of a target gene, as detailed below.

### Helenus feature (1): Kassandra deconvolution

Kassandra deconvolution predicts the RNA fractions of TME cell types (14), which was used as a feature to train the Helenus models in estimating the TME contribution to total gene expression of a sample. The Kassandra prediction consists of RNA percentages of different TME cell populations.

### Helenus feature (2): weighted average expression profile of the TME

Kassandra predictions were also used to estimate the TME contribution to the average expression profile. The RNA percentages of the TME cell populations were used as weights in the weighted sum of the average expression of a target gene in different TME cell populations. The average expression profile of one particular cell type is computed as the median expression of 18,792 genes across all pure samples of the cell type in the database:

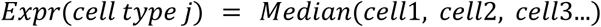

where *Expr*(*cell type j*) is the expression profile of the one cell type.

The average expression profile of the TME weighted by fraction of the TME cell types as estimated by Kassandra deconvolution can be used as a linear subtraction method for gene expression profile reconstruction or here, as an input for internal Helenus calculations:

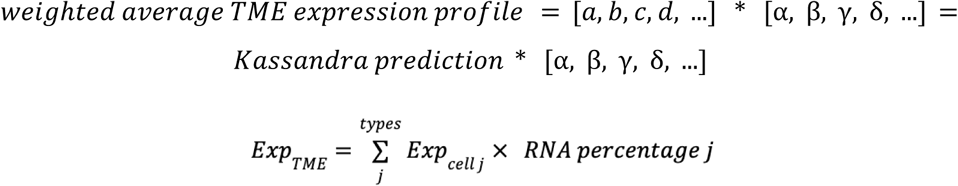

where [α, β, γ, δ, …] each corresponds to the average expression profile of a TME cell type and [*a*, *b*, *c*, *d*, …] each represents a vector of weights. The RNA percentages (weights in the weighted sum) of the different TME cell populations are predicted by Kassandra deconvolution.

### Helenus feature (3): gene selection

We employed genes used for training Kassandra models (> 20,002 genes) as the initial set of TME-specific genes. Genes with variable expression across major TME cell types in the samples used to model the TME were selected. The gene is considered as “variable” across TME cell types if:

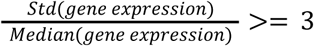

This method identified 475 TME-specific genes with variable expression. Additionally, the final set of genes included 127 clinically important malignant tumor biomarkers based on published literature, such as *EGFR* (36), *ERBB2* (37), and *CCND1* (38). Some models demonstrated low performance on validation data, and further optimization led to another approach of feature selection that is based on paired-occurence in pathways. We used the lists of genes that compose pathways from The Molecular Signatures Database (MSigDB, v7.5.1):

hallmark gene sets (h.all), curated gene sets - canonical pathways (C2.cp), computational gene sets - cancer gene neighborhoods (C4.cgn), oncogenic signature gene sets (C6.all), and immunologic signature gene sets (C7.all). For each target gene (i.e. model), we iterated all pathways in which the target gene is involved. Each gene that participates in the same pathways together with the target gene was scored +1. Then, we selected 600 genes with the highest score that corresponds to genes most often found in the same pathways as the target gene.

Models with improved estimates of MC expression (compared to the total expression profile) according to model criteria, used genes selected based on co-occurrence within pathways as features rather than those selected based on variability.

### Helenus model training parameters

Helenus is a multi-step ML approach for reconstructing the expression profiles of TME and MCs, developed using Python 3 with Pandas libraries for data manipulation and preprocessing (v1.5.3) (39,40), and NumPy for numerical and statistical operations (v1.22.4) (41,42).

Each model was trained to predict the weighted TME expression profile represented in the sample using LightGBM version 3.0.0 (43,44). The parameters used were: subsample: 0.9607; subsample_freq: 9; colsample_bytree: 0.2933; reg_alpha: 3.9006; reg_lambda: 2.938; learning_rate: 0.05; max_depth: 11; min_child_samples: 271; num_leaves: 2048; n_estimators: 3000.

### LightGBM hyperparameter optimization

LightGBM hyperparameters were selected separately for each gene model that did not pass criteria using the Optuna framework (v. 2.10.1). We optimized “boosting_type”, “max_depth”, “subsample”, “colsample_bytree”, “reg_alpha”, “reg_lambda”, “min_child_samples”, and “num_leaves”. For all models, we ran 500 trials with 1,000 trees in each ensemble with 0.05 learning rate. The performance of each combination of hyperparameters was validated using holdout artificial transcriptomes. Negative mean squared error was used as the objective function.

### Feature importance

The gene that the LightGBM model was trained for was expectedly among the 10 most important features for 1,068 out of 1,078 models incorporated in Helenus, as the expression of the gene in mixed samples can be used for rough estimations of gene expression in compartments (**Fig. S1C**). The weighted average expression profile of the TME reconstruction feature was among the 10 most important features for 712 out of 1,078 models, suggesting that weighted average expression profile was moderately beneficial for expression profile reconstruction. Other important features included gene expressions, often related to gene function or related processes, as well as deconvolution predictions of TME cell fractions.

For example, for the *CD274* (PD-L1) model, all genes serving as important features for model prediction were connected to immune system function or PD-L1 regulation directly, or co-regulated with PD-L1, such as *CD80* (45,46), *PDCD1LG2* (47,48), *IL27* (49), and *CCL22* (50), among others. For the *ERBB2* (HER2) model, important features for its expression profile reconstruction included genes expressed by fibroblasts and associated with epithelial to mesenchymal transition (EMT), like *MMP2* (51), *ROR2* (52), *CCL7* (53), *FGFR1* (54), *COL3A1*, and *COL1A2* (55). Interestingly, the percentage of fibroblast cells predicted by deconvolution was also an important feature. Noteworthily, *ERBB2* expression was not used by the deconvolution model to predict fibroblasts fraction; therefore, this feature was not merely due to an intersection between two algorithms (Helenus and deconvolution), and likely linked to relevant biological processes encountered by machine learning.

### Criteria of model selection

First, 1,885 models were trained to predict the expression profiles of several biomarker genes, genes involved in biochemical pathways, and widespread signatures. Then, 1,078 high-performing models that passed the selection criteria in validation tests were selected. The selection criteria were based on MAE/Mean reduction values pre- and post-Helenus (tumor compartment) application. The selected genes were split into different categories (TME-specific, MCs-specific, and shared genes) and then analyzed separately. The reference value included 133 samples of experimental data (*in vitro* mixes and sorted cells). A filtering step further excluded 260 genes from validation as their expression level was below the 5-TPM threshold in more than 5 samples. For all genes, the model was validated if MAE/Mean reduction was equal or more than twofold. TME-specific gene models were validated if both the predicted and real MC expression levels were below 5 TPM, while tumor-specific/shared gene models were validated if MAE/Mean values before and after Helenus application were less than 0.25 or MAE/Mean after Helenus application was less than 0.2).

#### Kaplan-Meier survival analysis and hazard ratios (HRs)

The Kaplan-Meier estimator was used to compare, over time, the estimated survival of patients with low, medium, and high biomarker expression levels pre- and post-Helenus application. The lifelines package (v0.28.0) (56,57) was used to plot the survival curves and calculate the *P*-values for the logrank test. The HRs and their 95% CIs were calculated using Cox proportional hazards regression models to assess the association between the gene expression values and the outcome.

### Quantification and statistical analysis

Statistics were calculated using the SciPy scipy.stats module in Python 3 (v1.8.1) (58). All graphs were plotted using a custom implementation of Matplotlib (v3.5.2) (59,60) and seaborn libraries of Python 3 (v0.11.2) (61). Scikit-learn was also used for statistical analysis and model evaluation (v1.1.1) (62,63).

The performance of reconstruction methods was evaluated with core metrics: (MAE/Mean) and concordance correlation coefficient (CCC). MAE/Mean metric was calculated to evaluate the mismatch between the reference and total (before reconstruction) or reconstructed gene expression profiles of MCs as:

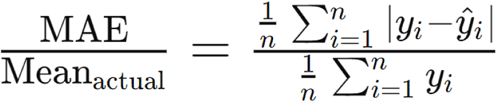

where y_i_ represents the actual value (true value) for the i-th observation, and ŷ_i_ represents the predicted value for the i-th observation.

The CCC was calculated as:

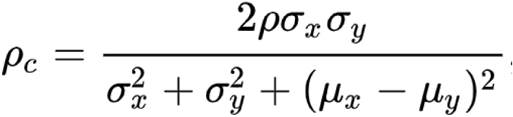

where ρ signifies the correlation coefficient between x and y, σ2 the variance and µ the mean. The Pearson correlation coefficients were also computed to compare Helenus-reconstructed gene expression profiles to other reconstruction methods, unless otherwise stated. The significance of Pearson correlations (r) to be non-zero was assessed by the use of the exact distribution of r (two-tailed test). Spearman correlation coefficients (ρ) were used to estimate correlations between scaled expressions and gene copy numbers; they were calculated with median values for each CNV value. Statistical comparison of two samples was conducted with Wilcoxon signed-rank test for paired samples (**Fig. 3C, E, G, I**) or Mann-Whitney U rank test for independent samples (**Fig. 4E-G**). Bonferroni correction for multiple comparisons was used.

Principal Component Analysis (PCA) was conducted with the sklearn.decomposition library fromscikit-learn (v1.1.1). Kaplan-Meier survival analysis was performed with the lifelines package (v.0.28.0). Logrank test was used to calculate *P*-values, and Cox proportional hazards regression models were used to calculate the hazard ratios (HRs) and their 95% CIs.

## ACKNOWLEDGMENTS

The results shown in Figure 4 and S3 are in part based upon data generated by the TCGA Research Network (phs000178): https://www.cancer.gov/tcga. This study also used publicly accessible data in the ArrayExpress, GEO, and the EGA (European Genome-Phenome Archive) (EGAS00001002928, EGAS00001004353), all cited in this study. Generative AI tools were used to refine the language and improve readability in certain parts of the manuscript. The authors have reviewed and edited the content as needed and take full responsibility for the content of this manuscript.

## DECLARATION OF INTERESTS

This research was funded by BostonGene Corporation. All authors affiliated with BostonGene, Corp. were employees thereof at the time the study was performed. A. Za., A. B., M. C., V. B., B. S., D. D., A. Zo., and M. S. are inventors on patents related to this work.

The authors declare no other competing financial interests.

## SUPPLEMENT

**Figure S1:**
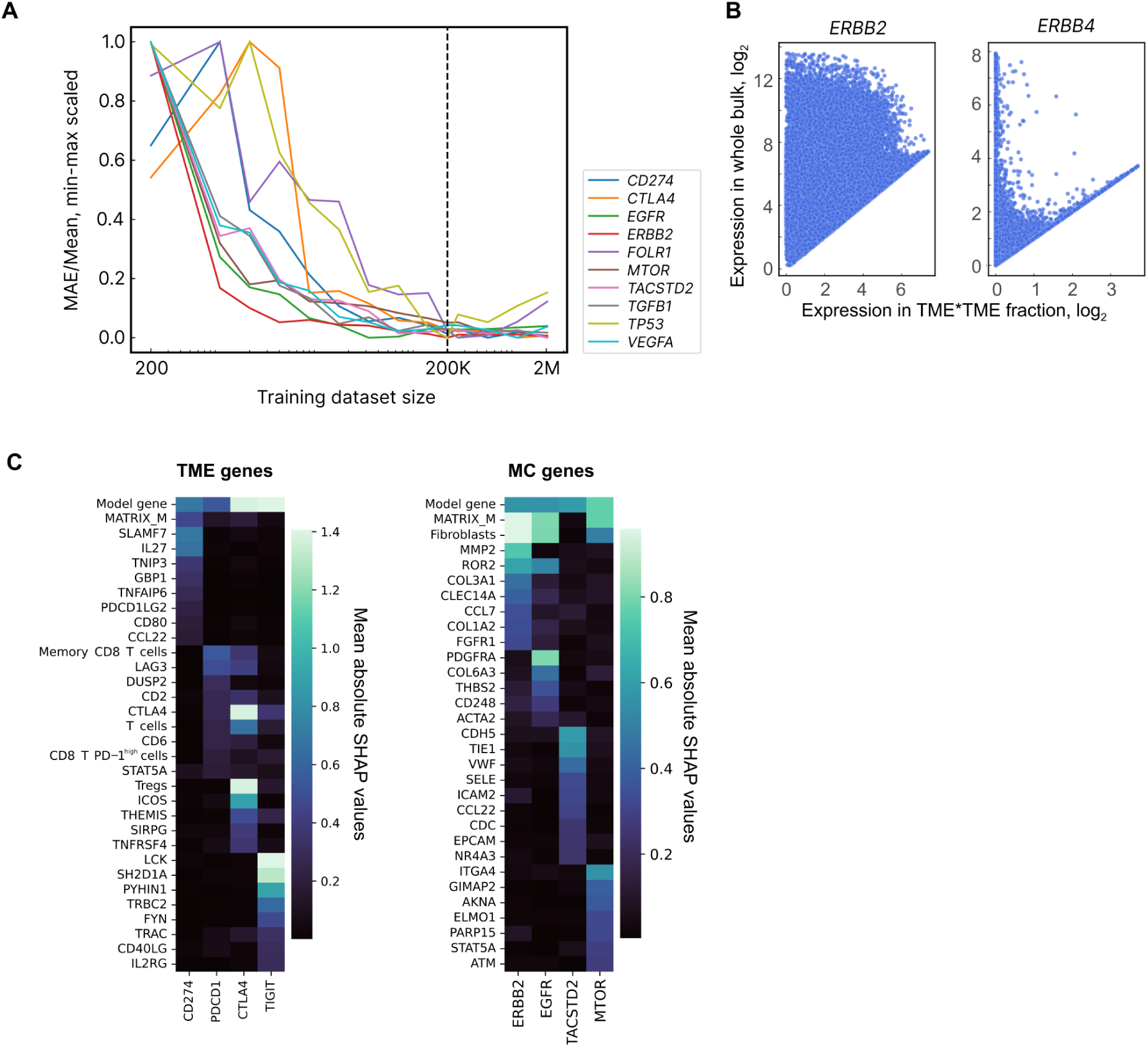
Evaluation of training dataset size and feature importance for Helenus development. Related to Fig. 1. **(A)** Training artificial transcriptome datasets of different sizes were evaluated with mean absolute error (MAE)/Mean values calculated on 248 validation samples for each model. Here, 10 models (one for one gene) tested across datasets with differing numbers of artificial transcriptomes are shown; the findings indicate that training datasets of 200,000 artificial transcriptomes per model are an optimal size. The MAE/Mean values were min-max scaled as models may differ in their performance, regardless of whether they were trained on datasets of the same size. **(B)** Among the 1,885 models generated using artificial transcriptomes as the training dataset for Helenus, 807 models failed to pass selection criteria (**Methods**). Bulk RNA-seq data for *ERBB4* as a function of its expression in the TME relative to the TME fraction offers a likely explanation: the limited expression coverage of *ERBB4* may have compromised the training and consequently, the performance of the model. In contrast, the *ERBB2* model passed the selection criteria, likely due to widespread expression of this gene in the samples of the training dataset. **(C)** The most important model features (color bar) for eight selected models are shown as examples: four models for TME-specific genes and 4 models for MC-specific genes. The 10 most important features (i.e., those with the highest mean absolute SHAP values) were selected for each model. SHAP values represent how much the feature affects model prediction. Since the effects may be positive or negative, absolute values were used. The gene for which the model was trained was among the most important features for all eight models shown here. Other important features included the Matrix M reconstruction result (’MATRIX_M’ feature for linear subtracting), expressions of genes, and deconvolution predictions for cell fractions such as for fibroblasts, memory CD8 T cells, and CD8 T PD-1^high^ cells.

**Figure S2:**
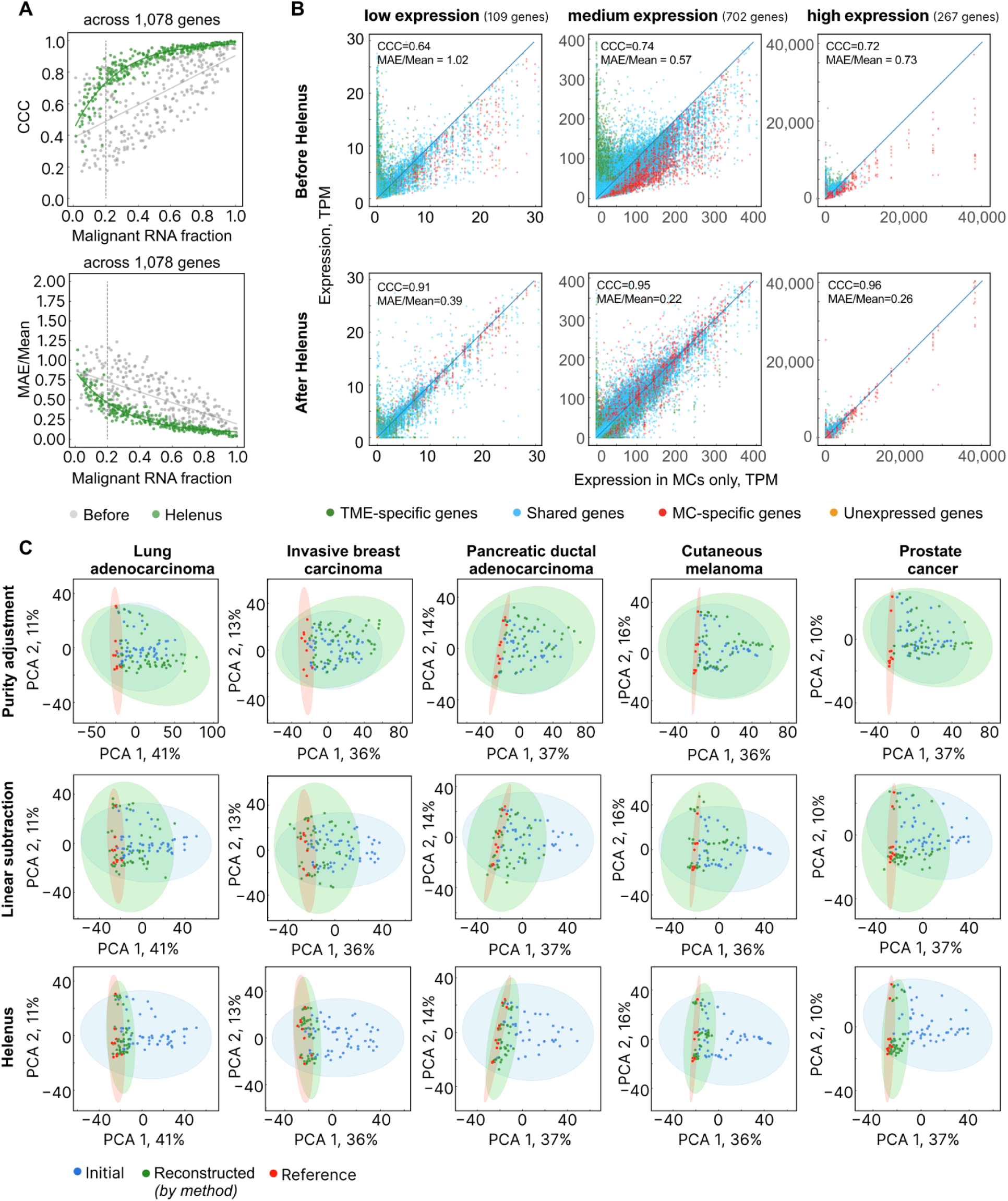
Gene expression reconstructions of artificial transcriptomes by Helenus simulating tumor purities, varieties, and expressions. Related to Fig. 2. **(A)** The similarity between MC profiles of artificial transcriptomes from holdout samples and those of reference MCs, as measured by pairwise concordance correlation coefficient (CCC) and the MAE/Mean, was limited for samples below the selected LOD of 20% tumor purity (dotted line). **(B)** Gene expression (TPM) in the mixed artificial transcriptomes before correction by Helenus were compared with the reconstructed expression profiles by Helenus for genes grouped by low, medium, and high average expression levels. Genes were also categorized as expressed in the TME (green), in MCs (red), in both theTME and MCs (blue), or as not expressed (orange). **(C)** Principal component analysis (PCA) projections for the five diagnoses separately to examine the impact of each tested approach (purity adjustment, linear subtraction, and Helenus, methods in green) on the whole expression profile of 1,078 genes, compared to initial expressions (blue) and reference (red). The PCA plots for holdout samples representing all five diagnoses together are presented in Fig. 2C.

**Figure S3:**
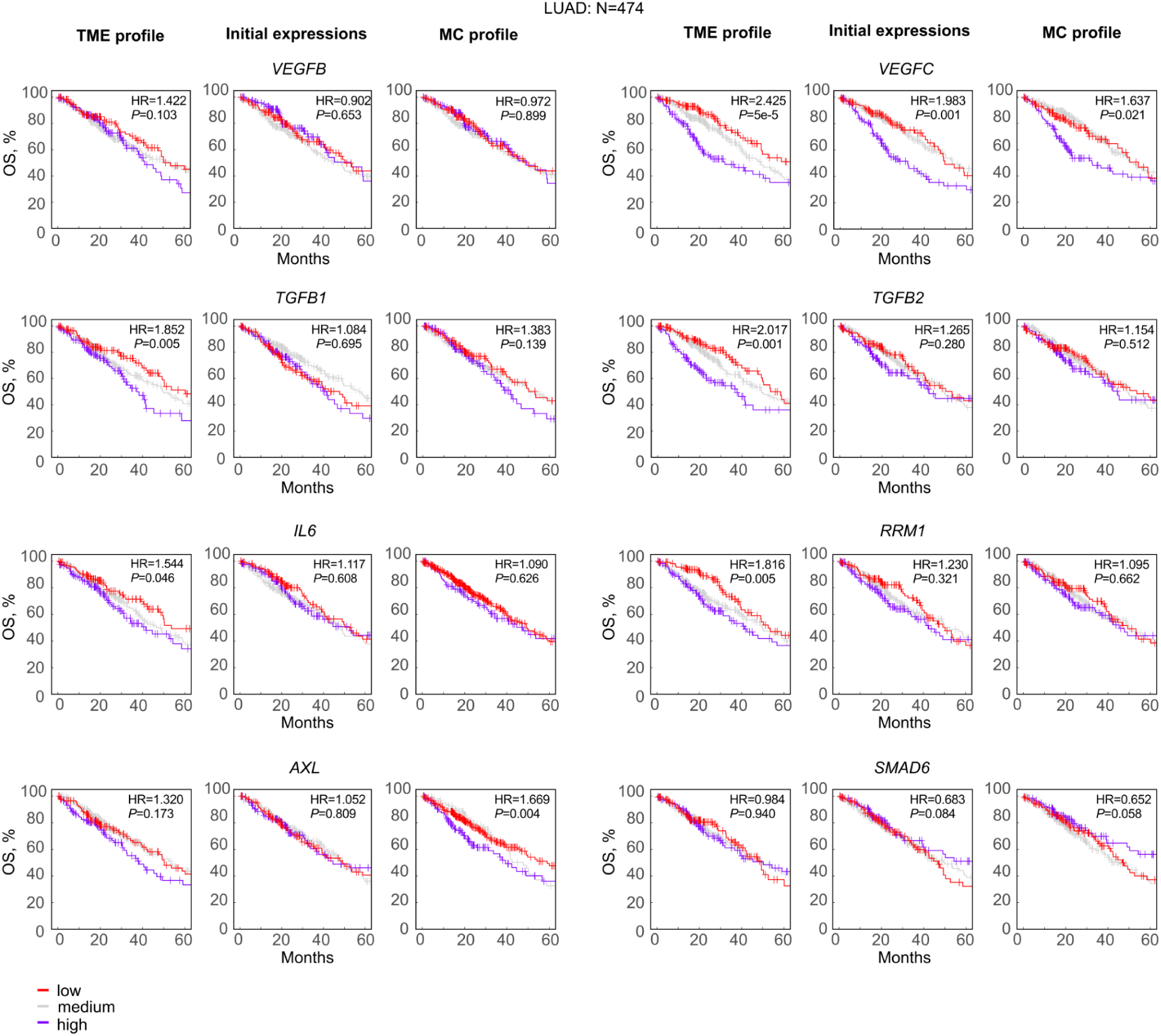
Survival analysis for patients stratified by corrected and uncorrected biomarker expression levels. Related to Fig. 4. Kaplain-Meier plots for TCGA-LUAD patients (N=474) based on uncorrected vs. Helenus-reconstructed expressions of individual biomarkers (logrank test). Patients were stratified into low (lower 25% quartile or 0q < x < 0.25q, red), medium (0.25q < x < 0.75q, gray), and high expression groups (upper 25% quartile or 0.75q < x < 1q, purple). Hazard ratios (HRs) indicate the survival differences between the high- and low-expression groups.

## Notes

### Competing Interest Statement

The authors have declared no competing interest.

